## Supplementary figures and images for "A novel machine learning algorithm selects proteome signature to specifically identify cancer exosomes"

### Supplementary Figures

Supplementary Figure S1

A

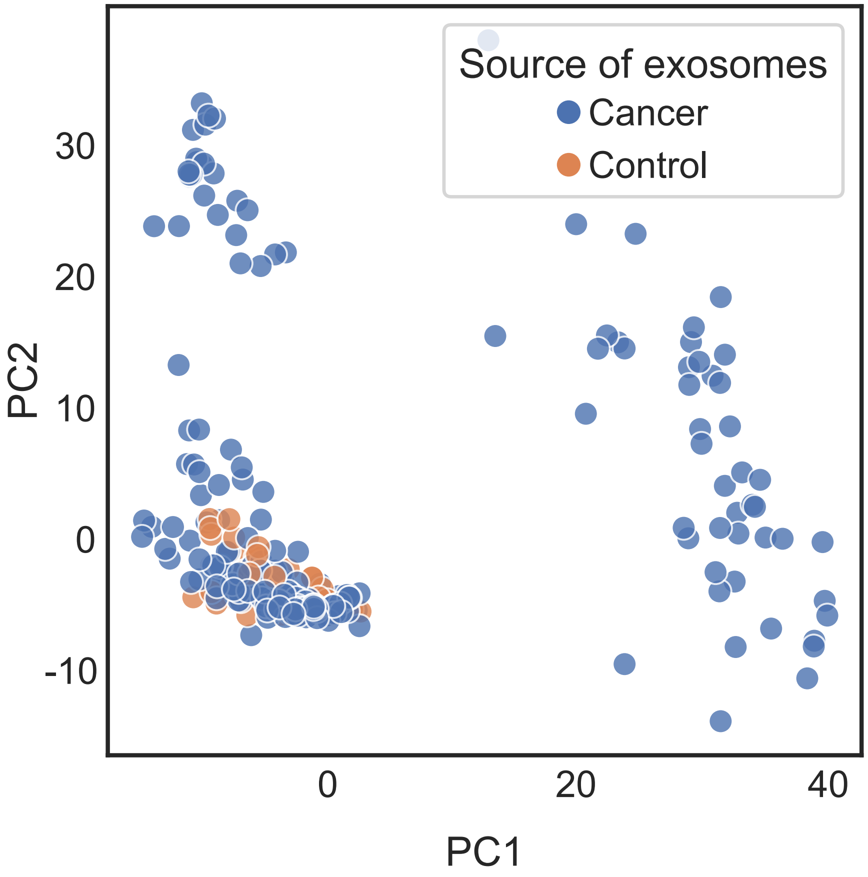


B

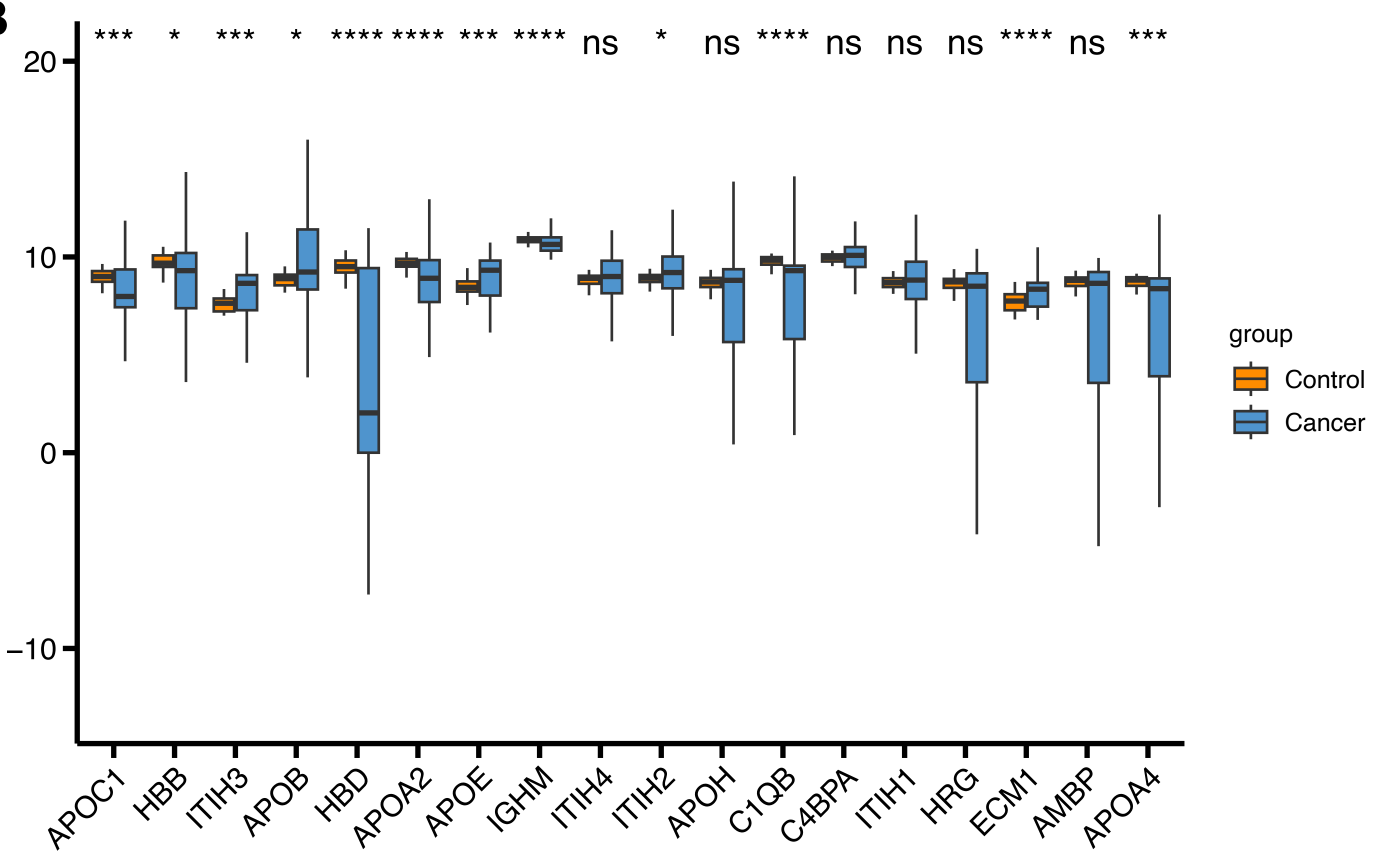
